# Demographic turnover undermines herd immunity in sympatric free-roaming dogs and cats: implications for zoonotic disease control in urban sentinel sites

**DOI:** 10.64898/2026.07.28.741324

**Authors:** Darren Norris, Fernanda Michalski

**Affiliations:** Ecology and Conservation of Amazonian Vertebrates Research Group, Federal University of Amapá, Macapá, Amapá, Brazil, 68903-419; Postgraduate Programme in Tropical Biodiversity, Federal University of Amapá, Macapá, Amapá, Brazil, 68903-419; School of Environmental Sciences, Federal University of Amapá, Macapá, Amapá, Brazil, 68903-419; Pro-Carnivores Institute, Atibaia, São Paulo, Brazil, 12945-010

**Keywords:** Demographic turnover, Vaccination coverage decay, One Health surveillance, Free-roaming dogs and cats, Zoonotic disease control, Sentinel Surveillance

## Abstract

Vaccination of free-roaming dogs (*Canis lupus familiaris*) and cats (*Felis catus*) remains a major public health challenge. Rapid urbanization forces these species into complex contact zones, where structural failure of pulsed vaccination under high demographic turnover undermines standard One Health interventions. In such cases species-specific intervention cycles are needed to reduce zoonotic disease risk. We integrated field-data within a simulated vaccination campaign to determine how species-specific turnover rates drive the erosion of herd immunity at an Amazonian urban sentinel site (university campus). We monitored free-roaming populations (72 dogs, 75 cats) using a non-invasive photographic mark-resight protocol from 2023 to 2025. We modelled time-to-disappearance using Cox Proportional Hazards and simulated the trajectory of effective vaccination coverage against a 40% herd immunity threshold, distinguishing between loss of vaccinated individuals and recruitment of susceptible individuals. The campus functioned as a high-turnover system, with 72% of dogs and 48% of cats classified as transients. Species significantly predicted persistence (Hazard Ratio = 0.56; 95% CI: 0.33–0.94; p=0.029), with cats exhibiting double the median residency of dogs (432 vs. 193 days). Consequently, the species experienced divergent epidemiological failure modes. For dogs, simulated vaccination coverage collapsed below a 40% herd immunity threshold in 160 days, driven by rapid immunity attrition (the loss of vaccinated individuals). Although cats persisted longer, their effective coverage was eroded by immunity dilution due to recruitment of naïve juveniles. Annual vaccination campaigns are likely insufficient in this high-turnover urban dog population. Effective One Health zoonotic control strategies must transition from static abundance-based targets to dynamic, species-specific and turnover-adjusted intervention schedules.

## 1. Introduction

Urban expansion creates contact zones among humans, domestic animals, and wildlife (Plowright et al., 2021). As human populations expand, land cover conversion into urban matrices exerts profound selection pressures on commensal and synanthropic species. Globally, this dynamic is uniquely volatile in tropical regions like the Amazon basin, where rapid, often disorderly urban expansion into sylvatic environments creates novel interfaces for zoonotic disease transmission (de Sousa Pereira et al., 2021, Lindahl and Magnusson, 2020, Hassell et al., 2017, Gibb et al., 2020, Ortiz-Prado et al., 2024). Within expanding urban environments, free-ranging domestic dogs (*Canis lupus familiaris*) and cats (*Felis catus*), occupy a unique and often precarious ecological niche (White and Razgour, 2020, Wilson et al., 2026, Langlois et al., 2025, Szentivanyi et al., 2024, Mendoza Roldan and Otranto, 2023, Viozzi et al., 2026). They act simultaneously as subsidized predators, dependent on human-provided resources, and as environmental sentinel hosts bridging the gap between endemic sylvatic pathogens and urban human populations (Combs et al., 2022, Nayeri et al., 2025). For example, in the Eastern Amazon, free-roaming dogs are vital sentinels for urban zoonoses, including Chagas disease and bat-mediated rabies (Bastos et al., 2021, Silva et al., 2024, Novaes et al., 2024).

Urbanization creates challenges for zoonotic disease control as it fosters large populations of free-roaming dogs and cats that are often too dispersed and mobile to monitor comprehensively. Therefore, effective surveillance requires logistically viable integrated approaches (Szentivanyi et al., 2024). Epidemiological monitoring can be strengthened by using sentinel sites - bounded, logistically tractable locations that are representative of the wider system where sustained, individual-level monitoring is feasible: (Guillot et al., 2022). Companion animals are increasingly recognized as sentinels within this framework. Because dogs and cats share housing, food, water, and parasites with the people around them, their infection patterns provide early warnings of zoonotic risk in the surrounding human population, a role documented for rabies and a range of other pathogens (Day et al., 2012, Kang et al., 2026). Resident free-roaming animals on university campuses occupy an analogous niche. Cared for by the campus community despite lacking formal ownership (Bicalho et al., 2024), they share the same exposure pathways as companion animals. However, campus populations span a dynamic range from these settled, community-provisioned residents to short-term transients.

University campuses meet several of the practical sentinel criteria for free-roaming dogs and cats (Guillot et al., 2022). Populations are spatially bounded and accessible, allowing animals to be photographed, and individually re-identified on a regular schedule. Critically, however, campuses are not closed populations. Free-roaming dog populations range from demographically stable, where pulsed vaccination remains effective, to high-turnover “sinks”, where losses and replacement quickly erode vaccine-derived immunity (Morters et al., 2014a, Morters et al., 2014b). Long-term, managed university cat colonies in Australia continually received immigrant or abandoned animals (Swarbrick and Rand, 2018), consistent with a “vacuum effect” where incoming animals colonize vacated territories (Miller et al., 2014). This continual pet abandonment is a well-documented challenge across universities, including in Brazil (Bicalho et al., 2024). Consequently, maintaining effective vaccination coverage of a campus is governed by population turnover, not merely abundance, and it is exactly this property that makes these sites informative sentinels.

Epidemiologically, the immune fraction of a vaccinated free-roaming population is set by which individuals remain present, not by how many animals are counted on any given day (Conan et al., 2015, Morters et al., 2014a). Yet most campus studies have focused on estimating abundance and demographic composition rather than tracking longitudinal residency (Bicalho et al., 2024, Phiriyaphokhai et al., 2025, Swarbrick and Rand, 2018). To date none have quantified individual-level residency or turnover rates — the parameters that ultimately determine whether pulsed vaccination coverage can persist between campaigns to maintain herd immunity.

Free-roaming dogs and cats occupy a critical position at the interface of human, domestic-animal, and wildlife health, acting as bridge hosts that sustain and transmit zoonotic pathogens across all three compartments. Beyond rabies — the paradigm pathogen motivating global vaccination targets — free-roaming dog and cat populations also maintain other viral diseases of veterinary and public-health concern (Chomel, 2014, Tomori and Oluwayelu, 2023), including Canine Parvovirus and Canine Distemper Virus, for which a comparable ∼70% herd-immunity target applies. Consequently, urban public spaces like university campuses, where free-roaming animals interact daily with dense, mobile human populations, function as key transmission nodes linking urban wildlife, domestic animals, and people.

Dogs and cats were domesticated through different evolutionary trajectories, which likely shape the demographic mechanisms driving vaccination outcomes. Dogs were actively selected for dependence and cooperative phenotypes, embedding them within human households and work (Kukekova et al., 2022, Lord et al., 2020, Pendleton et al., 2018). In contrast, cat domestication preserved their capacity for a solitary, self-sustaining existence. Even as they adjusted social structure, diet, and activity rhythms to human-shaped environments, cats retained the behavioral flexibility to persist independently of direct human provisioning or attachment (Natoli et al., 2022, Prigent Garcia and Chebly, 2024, Vitale, 2022, McGrath et al., 2024, Montague et al., 2014, Pongrácz and Lugosi, 2024). These contrasting behaviors provide a basis for the differing epidemiological patterns in urban areas. Additionally, this behavioral and cultural asymmetry extends directly to research investment itself. Dogs, celebrated across cultures as loyal companions, have attracted sustained veterinary and epidemiological attention, particularly for population management and rabies control. Cats, by contrast, are more often framed by conservation biologists as an invasive predator — ranked 38th among the world’s worst invasive species (Global Invasive Species Database, 2026). This divergence in cultural and disciplinary framing offers a likely explanation for why the vaccination-coverage data are relatively robust for dogs but remain comparatively scarce for cats.

Controlling zoonoses like rabies in free-roaming dogs and cats relies on sustaining a herd immunity threshold sufficient to interrupt transmission –defined by the World Health Organization as at least 70% initial coverage (Hampson et al., 2009). Critically, however, herd immunity built through pulsed mass vaccination decays continuously Modelling studies demonstrate that even when initial coverage reaches 60–80%, population-level protection can fall below critical thresholds within 12-24 months (Conan et al., 2015, Morters et al., 2014a, Morters et al., 2014b). This decay stems from a variety of factors including two distinct demographic processes that vaccination planning rarely separates. Vaccinated individuals are lost from the population through death or emigration, directly eroding the immune fraction — a process we term immunity attrition. Independently, unvaccinated individuals enter through birth or immigration, diluting the protected proportion of the population even where previously vaccinated animals remain present — a process we term immunity dilution. Because these mechanisms are governed by different aspects of a species’ life history, they are likely to respond to different management actions. Yet, few studies quantify their relative contributions, and to our knowledge, none compare them between sympatric species sharing a single environment.

Previous studies have documented demographic and turnover estimates of dogs in urbanizing or urban settings (Shamsaddini et al., 2022, Smith et al., 2022, Morters et al., 2014b, Belo et al., 2017). Studies have also examined demographic and abundance/density estimates for cats in urban settings (Cove et al., 2023, Gunther et al., 2020, Bennett et al., 2021, Flockhart et al., 2022). Across empirical, modeling, and review literature, high demographic turnover consistently undermines persistence of vaccination coverage in both species. Maintaining effective herd immunity therefore requires frequent, high-coverage campaigns paired with context-specific population management and monitoring appropriate to the local context (Conan et al., 2015, Chidumayo, 2018, Fehlner-Gardiner et al., 2024, Roebling et al., 2014).

Most One Health demographic and vaccination-decay models focus exclusively on dogs, reflecting the historical emphasis of rabies-elimination priorities (Belsare and Vanak, 2020, Chidumayo, 2018, Conan et al., 2015, Hudson et al., 2019a, Hudson et al., 2019b, Morters et al., 2014a, Morters et al., 2014b). Comparable demographic data for free-roaming cats within a zoonotic or public-health framing remain severely limited. Crucially, no study to our knowledge has evaluated how two distinct drivers of coverage decay – immunity attrition (loss of vaccinated individuals) and immunity dilution (recruitment of naïve individuals) – operate simultaneously and diverge between sympatric species.

To address these knowledge gaps, we evaluated the persistence of sympatric free-roaming cats and dogs and determined how this may influence herd immunity within an urban sentinel site (Marco Zero campus). We hypothesized that: (1) high population turnover rapidly erodes herd immunity, rendering standard annual vaccination pulses ineffective; and (2) sympatric dogs and cats exhibit divergent epidemiological failure modes. By modeling vaccination coverage decay to identify critical time points where protection falls below a 40% threshold, we illustrate how species-specific turnover parameters can guide dynamic, turnover adjusted One Health zoonotic control strategies.

## 2 Methods

### Ethics statement

This study was conducted in strict accordance with Brazilian regulations regarding animal research. The protocol involved purely observational, non-invasive data collection (visual surveys and photographic mark-resight) of free-roaming animals in public spaces. No animals were captured, handled, sedated, or manipulated, and the study did not involve any changes to the animals’ environment or diet. As the methodology did not involve “animal use” as defined by Brazilian Law 11.794 (Lei Arouca) and CONCEA Normative Resolution No. 30, formal approval from the Committee on Ethics in the Use of Animals (CEUA) was not required.

### 2.1 Study area

The study was conducted at the Marco Zero campus of the Universidade Federal do Amapá (UNIFAP), located in the municipality of Macapá, state of Amapá, northern Brazil (00°00′41″ N, 51°04′27″ W). The Marco Zero campus offers a highly exposed, resource-limited proxy for the broader municipal landscape, particularly because the institution does not have a veterinary medicine course to manage these abandoned populations internally. The campus sits directly on the Equator, covering a total area of approximately 0.93 km². The campus lies within a peri-urban landscape bordered by expanding residential neighborhoods and commercial areas. The campus is surrounded by neighborhoods of mixed socioeconomic status, where pet ownership is common but veterinary services and animal-control programs are limited. The campus is a public space interacting directly with surroundings marked by high residential density and varying levels of social vulnerability, factors that historically contribute to the pressure of animal abandonment on university grounds.

As the central administrative and academic hub of the university, the campus houses 84 buildings, supporting seven academic departments. The institutional community comprises approximately 8,013 enrolled students and a workforce of over 1,160 public servants, including 654 permanent professors and 508 technical-administrative staff. With several thousand students, staff, and visitors circulating daily through the campus during the academic year. The campus encompasses built infrastructure (classrooms, laboratories, administrative buildings, student housing, restaurants, sports facilities, and parking areas) interspersed with green spaces, including lawns, and small fragments of secondary Cerrado forest. Habitat types relevant to free-roaming carnivores include Urbanized zones: dense building complexes with food outlets, garbage collection points, and student housing, providing anthropogenic food resources. Green open spaces: lawns, athletic fields, and landscaped areas used for recreation. Forest fragments and shrubland: patches of secondary Amazon forest and regenerating vegetation offering shelter and denning sites. Riparian or drainage areas: seasonal wetlands and drainage channels with dense vegetation.

Unlike several Brazilian universities UNIFAP does not have any formal institutional recognition or management plans for the stray cats and dogs on any of its 4 campuses. Reports from students, staff, and local animal-welfare volunteers indicate that free-roaming dogs and cats have been present on the Marco Zero campus for decades, likely due to a combination of abandonment and uncontrolled reproduction in surrounding neighborhoods. Informal feeding sites and ad-hoc care by volunteers occur intermittently. In recent years, student groups and faculty have discussed implementing sterilization and vaccination initiatives, yet population-management programs have not yet been implemented.

### 2.2 Population monitoring

Free-roaming animals were defined as unconfined dogs or cats observed without physical restraint and not under direct human control at the time of sighting. To assess the population dynamics of free-roaming dogs and cats, we conducted a non-invasive observational study across the university campus from August 2023 to December 2025. We employed a standardized visual encounter survey design, utilizing existing walkways. Surveys were conducted monthly during the dry season and only during the academic term time to minimize bias associated with seasonal fluctuations in animal detectability and activity/movement patterns. Sampling occurred over five consecutive days per month, stratified by time of day to coincide with expected peaks in crepuscular activity and avoidance of elevated midday temperatures: morning sessions (08:30 – 11:30) and late afternoon sessions (16:00 – 18:15). Data collection was performed by two independent teams, each consisting of 2–3 trained observers. To reduce the potential for observer bias teams always included at least one observer who had been monitoring the animals since 2023 (Meunier et al., 2019). Teams simultaneously surveyed distinct campus sectors. Observers walked at an average speed of 2 km/h (SD ± 0.7), scanning for animals. Upon sighting an animal, observers recorded time, species, sex (if discernible), and apparent health status. A non-invasive photographic mark-resight technique was employed to identify individuals. High-resolution photographs were taken of each animal using mobile phones, focusing on unique natural markings (e.g., pelage pattern, scarring) to create an individual identification catalog.

### 2.3 Data analysis

#### 2.3.1 Relative abundance and recruitment

Here we use relative abundance (individuals sighted per kilometer walked) to establish relative changes through time rather than absolute population size. Consistent with established protocols for free-roaming dog and cat surveillance, we monitored relative abundance to detect demographic instability relevant to vaccination planning (Cunha Silva et al., 2025, Ghimire et al., 2025, Gunther et al., 2020). This approach assumes that standardized monitoring indices correlate positively with population density, providing a sufficient proxy for tracking temporal trends. Time series were modelled using Generalized Additive Model (GAM), where sightings per kilometer were explained by species and sex. We fitted GAMs with a Tweedie error distribution, including species, sex, and a smooth term for time. To evaluate these temporal dynamics, time series data were modeled separately for each species. We employed a two-step modeling framework to accurately deconvolve the overall directional trend from the non-linear temporal fluctuations, thereby avoiding the mathematical confounding (concurvity) that occurs when competing parametric and highly flexible smooth terms are fitted simultaneously.

First, to quantify the overarching trajectory of population change, we fitted a parametric Generalized Linear Model (GLM) using a continuous numeric time variable to extract the linear coefficient and its standard error. Second, to evaluate the non-linear complexity and cyclical variations in the sighting rates, we fitted a Generalized Additive Model (GAM) incorporating a penalized thin plate regression spline (s()) for time. The complexity of these non-linear variations was assessed using the Effective Degrees of Freedom (EDF) derived from the smooth term. For both the GLM and GAM frameworks, models were fitted using a Tweedie error distribution. The Tweedie distribution was selected as it appropriately accommodates the non-negative, right-skewed, and zero-inflated continuous nature of the response variable (individuals sighted per kilometer). All models were constructed and evaluated using the mgcv package in R. Statistical significance for both the linear trajectory and the non-linear smooth terms was set at α = 0.05.

To evaluate the interactive effects of species and sex on temporal abundance trends, we fitted a Generalized Additive Model (GAM) using the mgcv package in R, specifying a Tweedie error distribution to accommodate the right-skewed, non-negative continuous response variable (individuals sighted per kilometer). We defined a composite grouping factor for each demographic cohort (e.g., female dogs, male cats) and included it as a parametric term to test for baseline differences in relative abundance. To account for distinct non-linear temporal trajectories among these groups, we applied independent penalized thin plate regression splines for time, conditioned on each cohort. Finally, to control for unmeasured daily environmental variation and pseudoreplication arising from multiple observations recorded under identical conditions, the specific survey date was incorporated as a random intercept using a random effect spline, with the overall model optimized via Restricted Maximum Likelihood (REML).

To evaluate species-specific differences in the accumulation of new individuals over time, we employed a complementary approach. After excluding the original baseline cohort, we constructed a continuous daily timeline of new arrivals for each species. First, we fitted a Poisson Generalized Linear Model (GLM) to the daily arrival counts to test whether the overall rate of new individual recruitment differed significantly between dogs and cats. Second, we applied a two-sample Kolmogorov-Smirnov (KS) test to the numeric arrival dates to determine if the temporal distribution differed between the two populations.

#### 2.3.2 Residency and persistence

We defined transients as individuals sighted only once or over a period shorter than a species-specific resighting threshold. We analyzed the distribution of inter-sighting intervals (time elapsed between consecutive sightings of the same individual) for all animals observed on multiple occasions. The threshold for declaring an individual as transient was defined as the 95th percentile of the inter-sighting intervals (dogs: 47 days; cats: 68 days). Previous studies discuss how “transients” might just be resident dogs with low detectability (Tiwari et al., 2018). We therefore expected that if residents had low detectability, it might be possible that transients could be residents that were missed. To test whether residents had low detectability we compared the detection rate defined as the proportion of survey days when individuals were sighted.

To evaluate site fidelity, we modeled the time-to-disappearance using a Cox Proportional Hazards model. Persistence was defined as the duration (in days) between an individual’s first and last sighting. To evaluate the factors influencing the persistence of stray animals on the university campus, we analyzed the time-to-disappearance for each individual. The response variable was defined as the duration (in days) from the initial identification of an animal until its disappearance (permanent absence) from the study area. An absence event was defined as a confirmed death or unconfirmed disappearance from the study area (status = 1). Animals that remained present at the end of the study were right-censored (status = 0). Adopted individuals were excluded from this analysis so that positive human interventions did not artificially inflate the estimated environmental hazard risk.

We employed a semi-parametric Cox Proportional Hazards model to estimate the risk of loss associated with three covariates: species (dog vs. cat), sex (male vs. female), and age class at initial sighting (juvenile vs. adult). To maximize sample size and account for dynamic recruitment, we utilized an open cohort design. The study population included both the initial cohort identified at the start of the study and new individuals (immigrants or abandoned animals) identified during subsequent monthly surveys. To objectively define the disappearance event without relying on arbitrary fixed intervals, we used the previously calculated resighting threshold for each species. Individuals not resighted for a duration exceeding this species-specific threshold relative to the study end date were classified as lost (Status = 1) at the time of their last sighting. Individuals with a time-since-last-sighting below this threshold were considered right-censored (Status = 0).

For the Cox Proportional Hazards analysis, the time origin (t=0) was defined as the date of first identification for each animal. The survival time was calculated as the duration between the first and last sighting. Animals entering the study at later dates were treated as new independent observations starting at t=0 of their respective tenure on campus. The model results are reported as Hazard Ratios (HR) with 95% confidence intervals, where an HR > 1 indicates an increased risk of disappearing from the cohort relative to the reference group. The proportional hazards assumption was verified for all covariates using Schoenfeld residuals. Additionally, Kaplan-Meier survival curves were generated to visualize the persistence probabilities of the cohorts over time, and differences between groups were assessed using the Log-Rank test. Statistical significance was set at α = 0.05.

#### 2.3.3 Modeling the Erosion of Herd Immunity

To quantify the epidemiological impact of persistence on vaccination coverage, we developed a deterministic decay model projecting the stability of vaccine-induced herd immunity over time.

This model assumes demographic equilibrium (exit balanced by recruitment), providing a conservative baseline for vaccination planning. We utilized the species-specific Population Half-Life *T*_1/2_ derived from the Cox Proportional Hazard model. The decay rate constant for each species was calculated as:

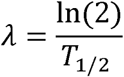

Where *T*_1/2_ represents the time (in days) required for 50% of the resident population to disappear from the study area. This metric serves as a proxy for the rate at which vaccine protection is lost from the specific focal population, independent of antibody titer waning.

We simulated the trajectory of effective vaccination coverage (*V*_t_) over a 12-month period (365 days) following a hypothetical mass vaccination campaign. We distinguish between the recommended initial campaign coverage (70%) and a lower functional herd immunity threshold (40%), below which transmission control is unlikely (Hampson et al., 2009). The model assumed an initial successful coverage (*V*_0_) of 70%, in accordance with World Health Organization (WHO) recommendations for eliminating dog-mediated rabies. The proportion of the population remaining vaccinated at time *t* was modeled using a standard exponential decay function:

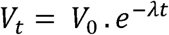

This model operates under the conservative assumption that individuals exiting the population are replaced by individuals entering the population (unvaccinated/naïve) at an equivalent rate, thereby maintaining a stable population size while diluting herd immunity. We identified the “Time to Critical Failure” for each species, defined as the number of days post-campaign (*t*_crit_) at which coverage drops below the herd immunity threshold of 40% (*V_crit_*). This was solved algebraically as:

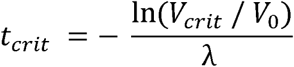

If *t_crit_* < 365 days, the population was classified as an unstable reservoir where annual pulsed vaccination strategies are mathematically insufficient to maintain continuous herd immunity.

#### 2.3.4 Observed Immunity Attrition and Dilution

We compared the theoretical decay models against empirical observation metrics to assess the impact of population turnover. To separate individual persistence from population-level immunity, we quantified two complementary trajectories: cohort retention and effective vaccination coverage. Cohort retention measures the proportion of individuals assumed to be vaccinated at the start of the study that remained present over time and therefore captures immunity loss due solely to death or emigration (immunity attrition). In contrast, effective vaccination coverage represents the proportion of the total population that remained protected at each time point, accounting for both the loss of vaccinated individuals and the recruitment of unvaccinated animals through birth or immigration. Divergence between these trajectories provides a direct measure of immunity dilution, where herd immunity declines despite the persistence of vaccinated individuals.

We calculated cohort retention *R_obs_*(*t*) as the proportion of the original cohort remaining on site at time *t*. This metric tracks the persistence of the individuals assumed to be vaccinated at *t*_0_:

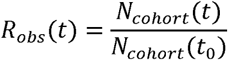

Where *N_cohort_*(*t*)is the number of original cohort members confirmed present on day *t*.

To measure Effective Vaccination Coverage *V_eff_*(*t*) we accounted for the influx of new, unvaccinated individuals (recruitment via birth or immigration). We calculated the daily proportion of the original cohort, adjusted by the initial vaccination coverage efficiency (*V*_0_):

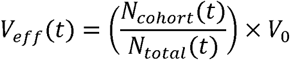

Where *N_total_*(*t*) represents the total census population on day (survivors plus new recruits).

We set the initial coverage (*V*_0_) at 70%, consistent with WHO recommendations for rabies elimination. We determined the “Time to Critical Failure” as the day on which *V_eff_*(*t*) dropped below the herd immunity threshold of 40%.

To explicitly quantify the relative contribution of demographic turnover mechanisms to the failure of herd immunity, we decomposed the total vaccination coverage loss into two distinct fractions: Immunity Attrition (loss of vaccinated individuals via death/emigration) and Immunity Dilution due to Recruitment (reduction in coverage via recruitment of naïve individuals).

We defined Total Coverage Loss Δ*V_total_* at time *t* as the difference between the initial target coverage (*V*_0_) and the observed Effective Vaccination Coverage *V_eff_*(*t*):

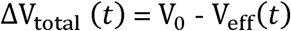

We then calculated the Attributable Fraction of Attrition Δ*V_attrittion_*, representing the loss of herd immunity caused solely by the death or emigration of vaccinated individuals. This was derived by calculating the theoretical coverage that would remain if no new individuals entered the population (i.e., assuming the population was a closed cohort):

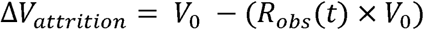

Whe *R_obs_*(*t*)is the cohort retention rate defined previously.

Finally, we calculated the Attributable Fraction of Dilution Δ*V_dilution_*, representing the additional reduction in coverage caused by the recruitment of naïve individuals (births or immigration) into the population. This was calculated as the difference between the theoretical closed-cohort coverage and the actual effective coverage:

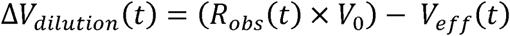

We converted these absolute values into percentage contributions to determine the primary driver of coverage failure for each species at the end of the study period (*t* = 365). A higher proportion of Δ*V_attrittion_* indicates that intervention failure is driven by the loss of vaccinated animals, while a higher proportion of Δ*V_dilution_* indicates failure driven by the influx of susceptible animals.

#### 2.3.5. Bootstrapped Estimation of Sample Confidence Intervals

To quantify the uncertainty associated with immunity attrition and demographic dilution without assuming a parametric distribution for residency times, we employed a non-parametric stratified resampling bootstrap approach. This method generates confidence intervals by accounting for the individual variability in residency and recruitment observed within the study population. The resampling procedure was stratified by species to preserve the original sample sizes (N_dog_ and N_cat_) and species-specific variances. For each of 1,000 iterations, we performed the following steps. Firstly, individuals were selected with replacement from the original dataset to create a “virtual” population of size. This process randomizes the composition of the cohort while maintaining the empirical distribution of residency durations and arrival times. Then, a daily presence-absence matrix was generated for the resampled population. Individuals present on the first day of the campaign were designated as the “Vaccinated Cohort” for that specific iteration. For every day of the study period, we then calculated Observed Attrition as the proportion of the specific iteration’s vaccinated cohort remaining on site(*N _survivors_*(*t*)/*N_cohort_*(*t*_0_)) and Effective Coverage as the proportion of the total virtual population consisting of vaccinated survivors ([*N_survivors_*(*t*)/*N_total_*(*t*)]×*V*_0_), where *V*_0_ represents an initial campaign coverage of 70%. Finally, the 95% Confidence Intervals (CI) for both metrics were derived from the 2.5th and 97.5th percentiles of the distribution generated across all iterations.

## 3. Results

### 3.1. Survey effort, cohort composition, and turnover

Between August 2023 and December 2025, surveys were conducted on 81 sampling days, covering a total distance of 310.9 km (Table 1). A total of 147 unique free-roaming individuals were identified, comprising 72 dogs and 75 cats. The initial cohort consisted of 45 individuals (28 dogs, 17 cats). By the end of the study period 18% of this cohort remained on campus (8 individuals; 4 dogs and 4 cats). Of those that left, 12 were adopted, 8 were confirmed dead on or within 15 m of the campus, and 16 disappeared. The final cohort consisted of 49 individuals (20 dogs, 29 cats). Across the study period, 102 new individuals were identified after the initial sampling phase, corresponding to a recruitment ratio of 2.3 new individuals per initial resident.

**Table 1.**
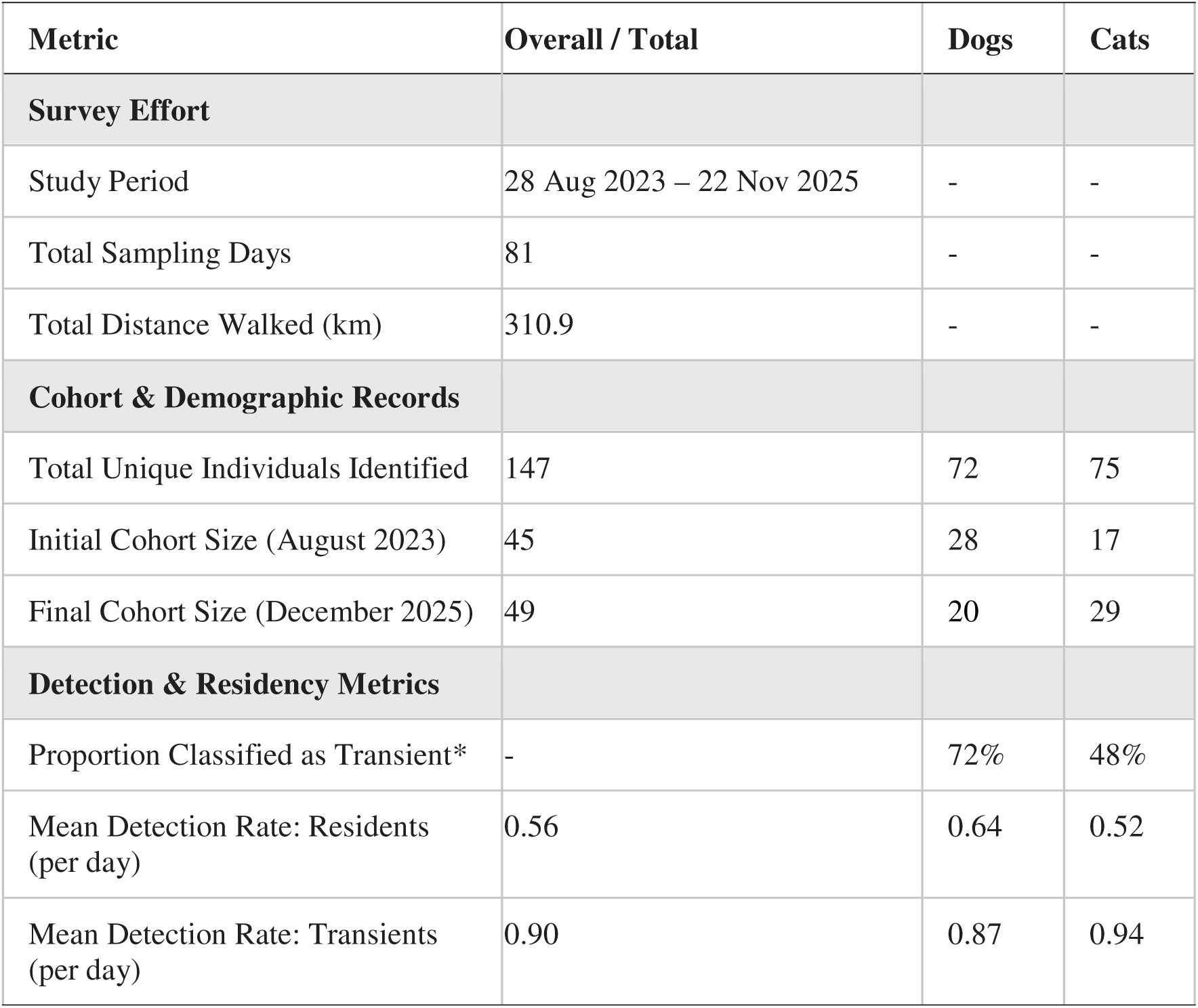
Summary of effort and records. Summary of survey effort, cohort demographics, and detection metrics for free-roaming dogs and cats on the Marco Zero campus. Transients are defined as unadopted individuals with residency durations below the 95th percentile of inter-sighting intervals (45 days for dogs; 68 days for cats).

Population turnover was high in both species. Using species-specific resighting thresholds (dogs: 47 days; cats: 68 days), 72% of dogs and 48% of cats were classified as transients (unadopted individuals with residency durations below the threshold). Consequently, only 42% of all identified individuals met the residency criteria. On average, resident animals were each recorded on 29 occasions, with a mean detection rate of 0.56 per day i.e. seen 50% of the time (mean detection rate of 0.64 and 0.52 for dogs and cats respectively). In comparison, transients were recorded on average 3 times, with a mean detection rate of 0.90 per day (mean detection rate of 0.87 and 0.94 for dogs and cats respectively). Overall, 37% (n=18) of transients were seen only once (42% and 30% of transient dogs and cats respectively).

### 3.2. Abundance trends and recruitment

The overall distribution of cat and dog abundances was similar (Figure 2a). The number of individuals sighted per km changed significantly over time (Figure 2b). Generalized additive models indicated species-specific temporal trends, with sighting rates of dogs declining over the study period, while sighting rates of cats increased (Table 2a). Dog sighting rates declined by 56.5% over the study period, while cat sightings increased by 122.2%. These temporal dynamics were non-linear and significantly structured by sex (Table 2b). Baseline sighting rates differed across all demographic cohorts, with female cats representing the most frequently observed group. Furthermore, while female cats, female dogs, and male dogs all exhibited significant temporal fluctuations in abundance, the relative abundance of male cats remained consistently lowest and statistically stable over time (Table 2b, Supplemental Material S1).

**Figure 1:**
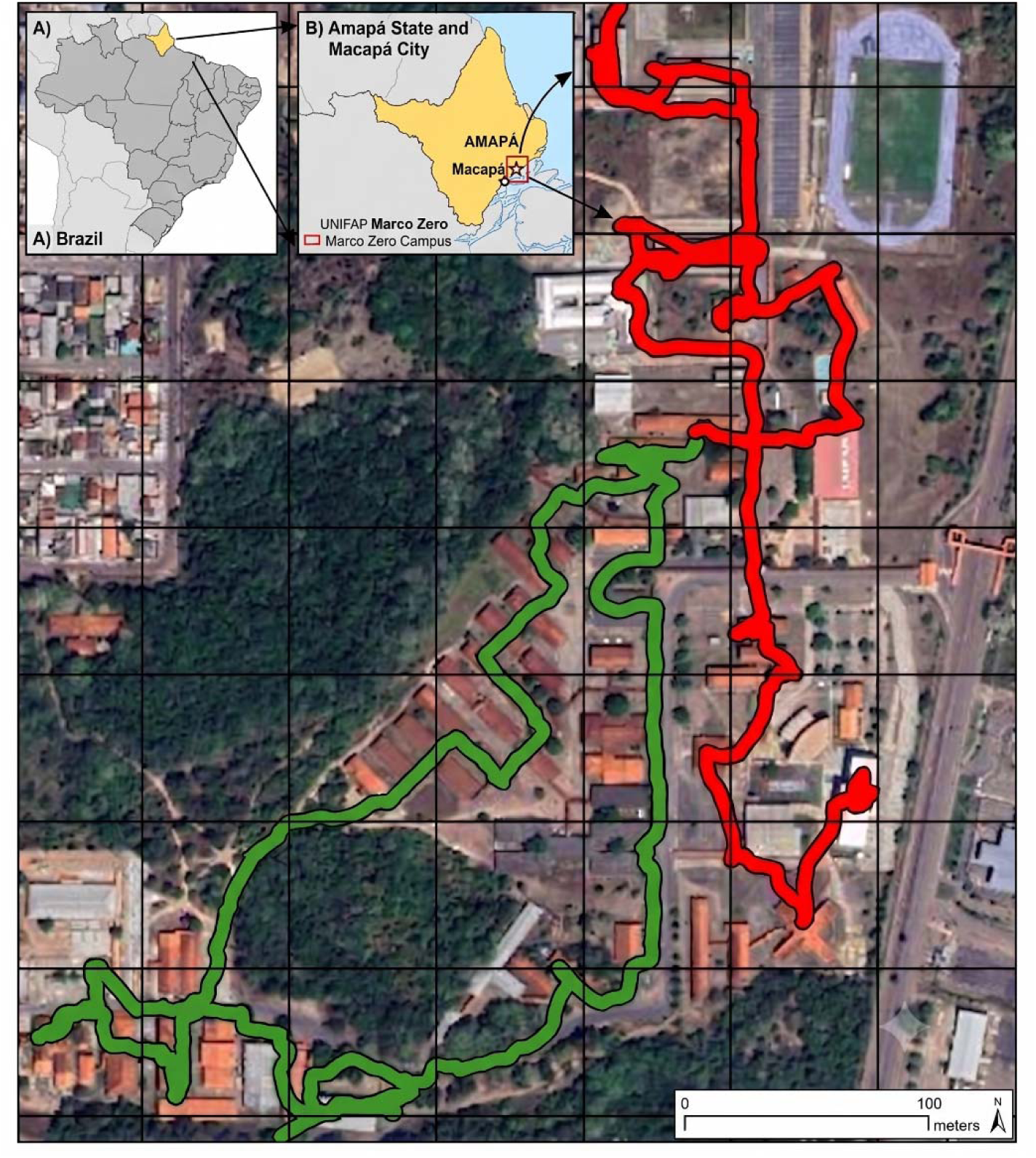
Study area. Survey location at the Marco Zero campus, Federal University of Amapá (UNIFAP), Brazil. The map shows the pathways used for standardized visual encounter surveys. All surveys were conducted along fixed routes within the campus perimeter. Cartographic Base: Google Earth v. 7.3.7.1155. Imagery provided by Maxar Technologies, Airbus (Image Date: November 17, 2025).

**Figure 2:**
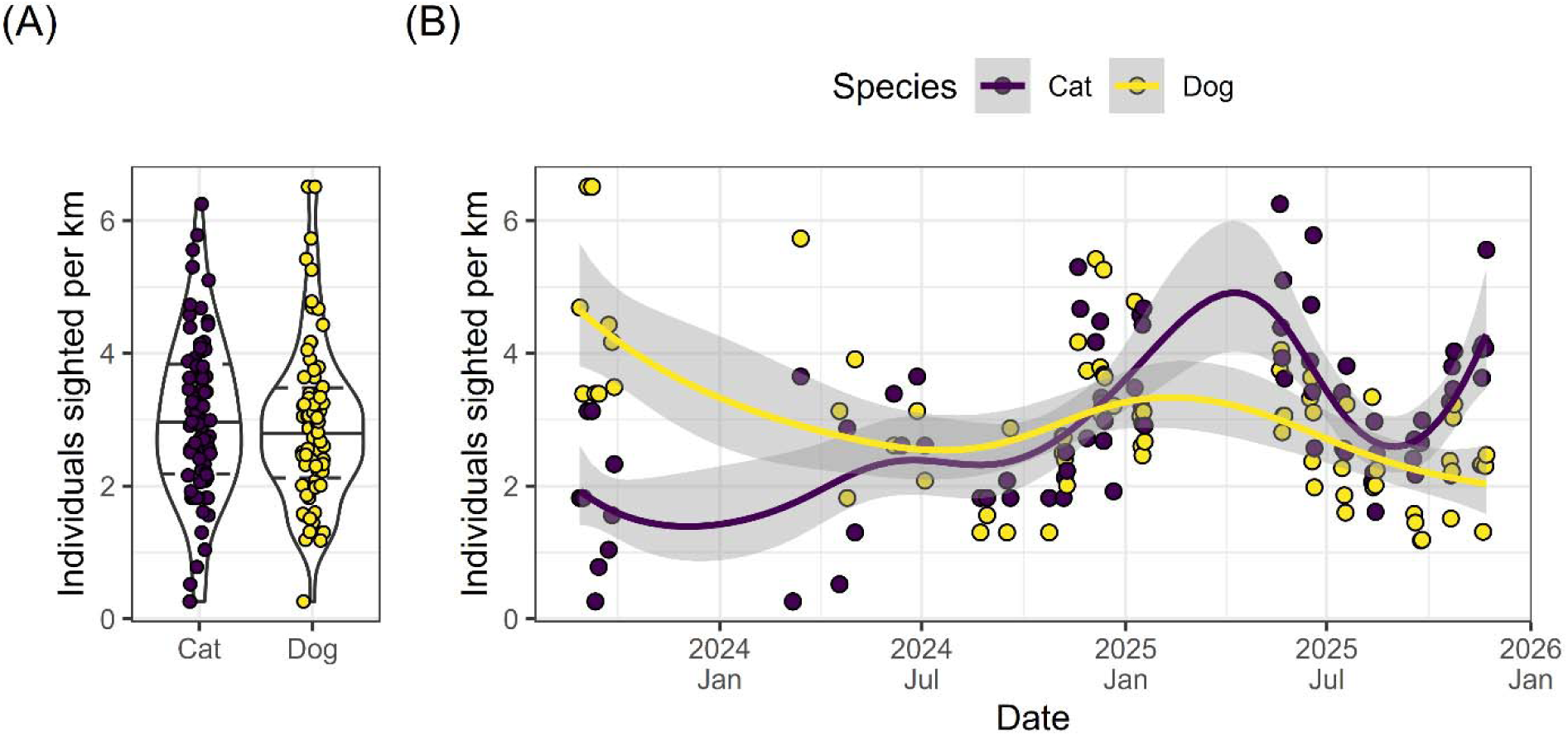
Abundance Trends. Relative abundance of free-roaming dogs and cats on the Marco Zero campus. Points are daily values, expressed as the number of individuals sighted per kilometer walked. (A) Distribution of daily sighting rates for dogs and cats across the study period. Violin plots show the density of observations; horizontal solid lines indicate medians and dashed lines indicate the 25th and 75th percentiles. (B) Temporal trends in relative abundance over time. Solid lines show predictions from Generalized Additive Models fitted separately by species to aid visual interpretation. Grey shaded areas indicate 95% confidence intervals.

**Table 2.** Summary of Temporal Trends (Tweedie GLM and GAM). (A) Parametric linear coefficients representing overall trend direction, alongside non-linear smooth complexities. (B) Summary of Generalized Additive Model (GAM) incorporating species, sex, and temporal terms.

| Species | Linear Coef | Linear SE | Linear p-value | EDF | Smooth F-statistic | Smooth p-value | GAM Deviance Explained (%) |
| --- | --- | --- | --- | --- | --- | --- | --- |
| Dog | -0.0007 | 0.0001 | < 0.001 | 4.1287 | 7.5660 | < 0.001 | 34.6 |
| Cat | 0.0008 | 0.0002 | < 0.001 | 6.8238 | 8.4042 | < 0.001 | 46.7 |

2b
| Model Term | Estimate / EDF | Std. Error | Test Statistic | p-value |
| --- | --- | --- | --- | --- |
| **Term Type: Parametric Terms** |  |  |  |  |
| (Intercept) | 0.643 | 0.036 | 17.733 | < 0.001 |
| Cat male | -0.976 | 0.046 | -21.276 | < 0.001 |
| Dog female | -0.230 | 0.037 | -6.270 | < 0.001 |
| Dog male | -0.443 | 0.039 | -11.451 | < 0.001 |
| **Term Type: Smooth Terms (Temporal Trends and Daily Random Effect)** |  |  |  |  |
| Time Trend: Cat female | 7.145 |  | 15.902 | < 0.001 |
| Time Trend: Cat male | 1.752 |  | 0.819 | 0.437 |
| Time Trend: Dog female | 5.848 |  | 8.725 | < 0.001 |
| Time Trend: Dog male | 3.955 |  | 7.287 | < 0.001 |
| Random Effect: Survey Day | 57.413 |  | 3.260 | < 0.001 |

The total number of new recruits identified over the study period was similar between the two species, with 44 new dogs and 58 new cats (Supplemental Material S2). There was no significant difference in the daily rate of new arrivals between cats and dogs (Poisson GLM, *p* = 0.167). The recruitment pressure per day was statistically equal across species. The temporal shape of the accumulation curves does not differ significantly (Kolmogorov-Smirnov test, D = 0.159, *p* = 0.414). This suggests that both species accumulated new individuals at a structurally similar rhythm over the survey timeline.

### 3.3. Individual persistence and drivers of disappearance

Persistence differed significantly between species. Cox proportional hazards analysis identified species as a significant predictor of time to disappearance (model Likelihood Ratio Test = 6.2, *p* = 0.045, Table 3). Cats exhibited a lower hazard of disappearance than dogs (HR = 0.56; 95% CI: 0.33–0.94; *p* = 0.029), corresponding to a 44% reduction in disappearance risk relative to dogs (Figure 3). Sex was not a significant predictor of persistence. Males exhibited a higher hazard of disappearance than females (HR = 1.27; 95% CI: 0.75–2.13), but this effect was not statistically significant (*p* = 0.37). No strong interaction between species and sex was detected (HR = 0.82; 95% CI: 0.25–2.67; *p* = 0.741). Median persistence time was more than twice as long for cats (432 days) as for dogs (193 days). This means that (adjusting for covariates), an individual cat had a 50% probability of surviving past day 432. Although males tended to persist for shorter durations than females, Kaplan–Meier survival curves reflected the similar patterns across sexes (Figure 3, HR = 1.27; 95% CI: 0.75–2.13; *p* = 0.369).

**Figure 3.**
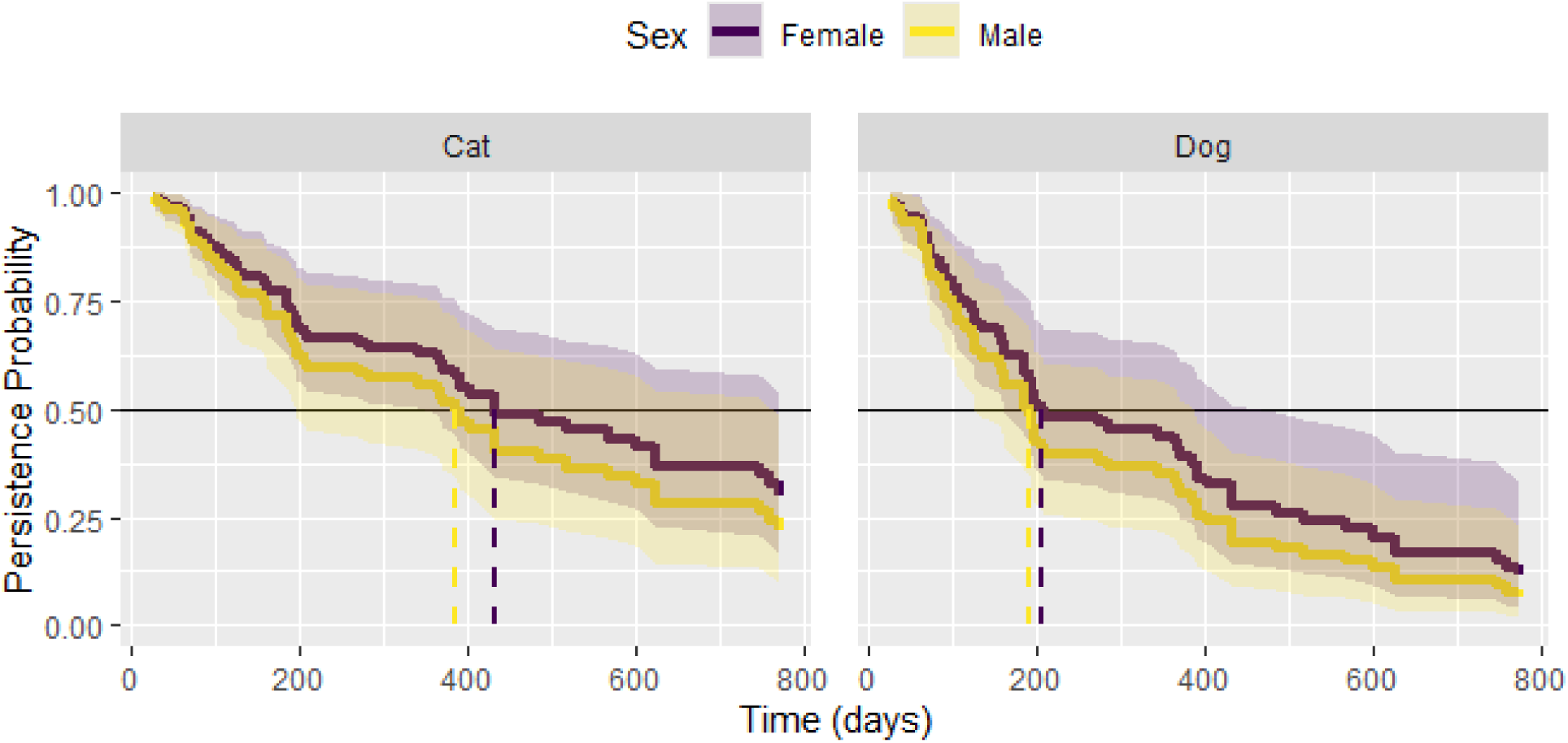
Persistence Probability. Persistence probability of unadopted free-roaming dogs and cats on the Marco Zero campus. Results of Cox Proportional Hazard model showing the probability of individuals remaining present on campus over time since first sighting. Solid lines represent the estimated probability over time for males and females. Shaded regions indicate 95% confidence intervals. Dashed vertical lines mark the median time for each group.

**Table 3.** Cox Proportional Hazards. Cox Proportional Hazards model results for unadopted dogs and cats at the Marco Zero Campus. Demographic breakdown of free-roaming dogs and cats monitored on the Marco Zero campus, reporting total sample sizes, events (confirmed death or unconfirmed disappearance), and right-censored outcomes (present at end of study), alongside Cox Proportional Hazards model estimates predicting time-to-disappearance. The event denotes a confirmed death or unconfirmed loss from the campus (Status = 1); censored denotes an individual still present on campus at the end of the monitoring threshold (Status = 0). Hazard Ratio (HR) estimates >1 indicate a higher relative risk of disappearance. Significant values (alpha ≤ 0.05) are marked with an asterisk. Overall Model Fit: Concordance Index = 0.625; Likelihood Ratio Test = 6.20 (*p* = 0.045).

| Demographic Group | N | Events (%) | Censored (%) | Hazard Ratio (95% CI) | p-value |
| --- | --- | --- | --- | --- | --- |
| **Overall Cohort** | **85** | **58 (68.2%)** | **27 (31.8%)** |  |  |
| **Species** |  |  |  |  |  |
| Dog (*Canis l. familiaris*) | 44 | 33 (75.0%) | 11 (25.0%) | Reference | - |
| Cat (*Felis catus*) | 41 | 25 (61.0%) | 16 (39.0%) | 0.56 (0.33 - 0.94) | 0.029* |
| **Sex** |  |  |  |  |  |
| Female | 46 | 29 (63.0%) | 17 (37.0%) | Reference | - |
| Male | 39 | 29 (74.4%) | 10 (25.6%) | 1.27 (0.75 - 2.13) | 0.369 |

### 3.4. Modeled erosion of vaccination coverage

Using species-specific population half-lives derived from the Cox model, simulated vaccination coverage declined rapidly in both species following a hypothetical mass vaccination campaign with 70% initial coverage (Figure 4). For dogs, modeled coverage fell below the 40% herd immunity threshold at 160 days post-vaccination. For cats, modeled coverage remained above this threshold until approximately 350 days. These projections reflect loss of vaccinated individuals under the assumption of demographic equilibrium, where exiting individuals are replaced by unvaccinated recruits at equivalent rates.

**Figure 4.**
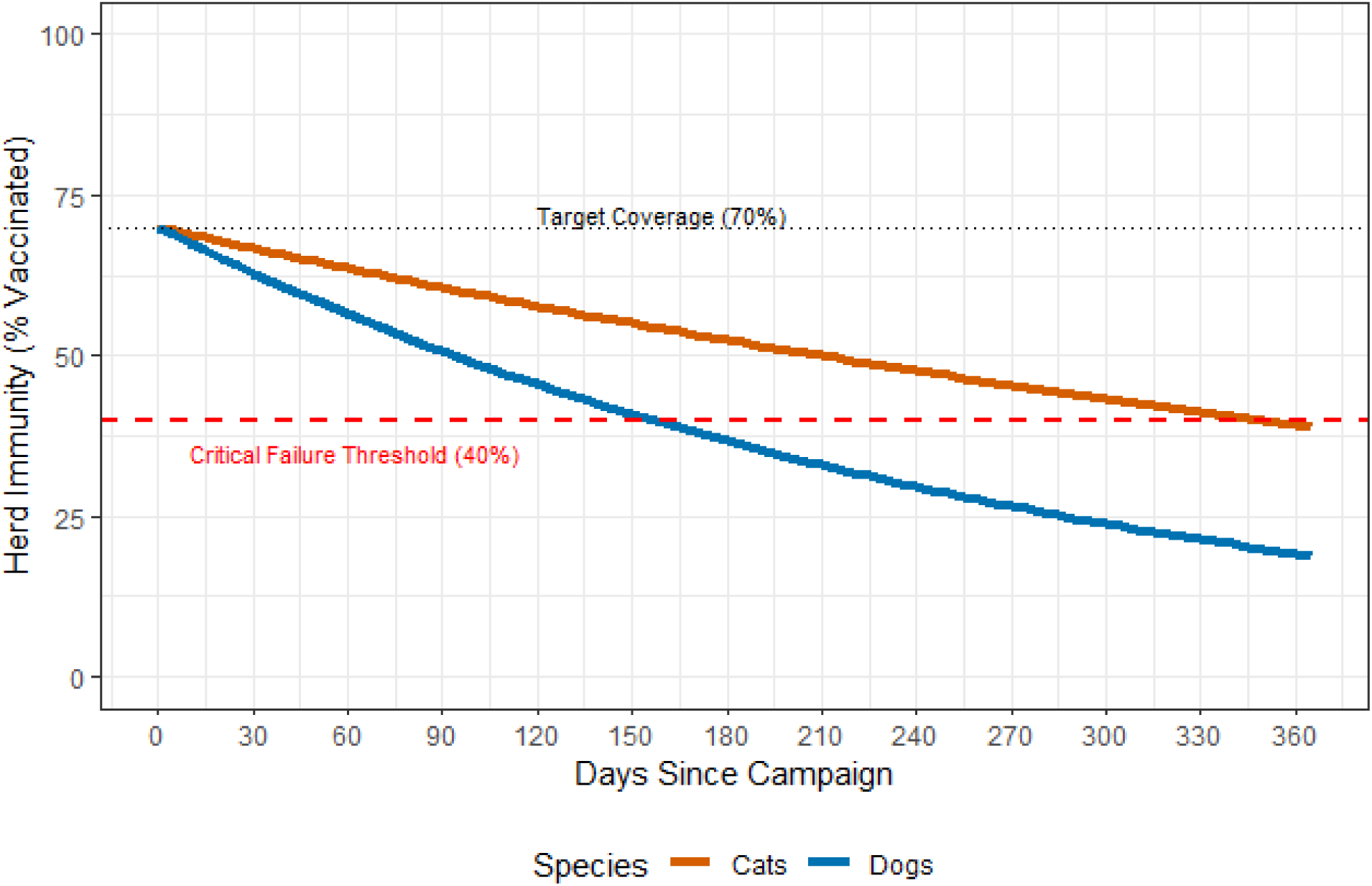
Modeled erosion of vaccination coverage. Modeled decay of vaccination coverage following a hypothetical mass vaccination campaign with 70% initial coverage of unadopted free-roaming dogs and cats on the Marco Zero campus. The solid lines illustrate the decay of herd immunity from an initial target of 70% (dotted black line) over a one-year period. Coverage trajectories were derived using species-specific population half-lives estimated from Cox proportional hazards models. The horizontal dashed red line indicates the herd immunity threshold of 40%. Vertical dashed lines mark the time at which modeled coverage falls below the threshold.

### 3.5. Observed immunity attrition and immunity dilution

Based on point estimates, empirical observations suggested a divergence between the persistence of vaccinated individuals and population-level immunity coverage (Figure 5). However, bootstrapped 95% confidence intervals revealed high individual variability, indicating that the strength of this divergence is species-specific and highly volatile. In dogs, effective vaccination coverage declined below the 40% threshold at 248 days, despite declining relative abundance. In cats, effective coverage declined below the threshold at 436 days.

**Figure 5.**
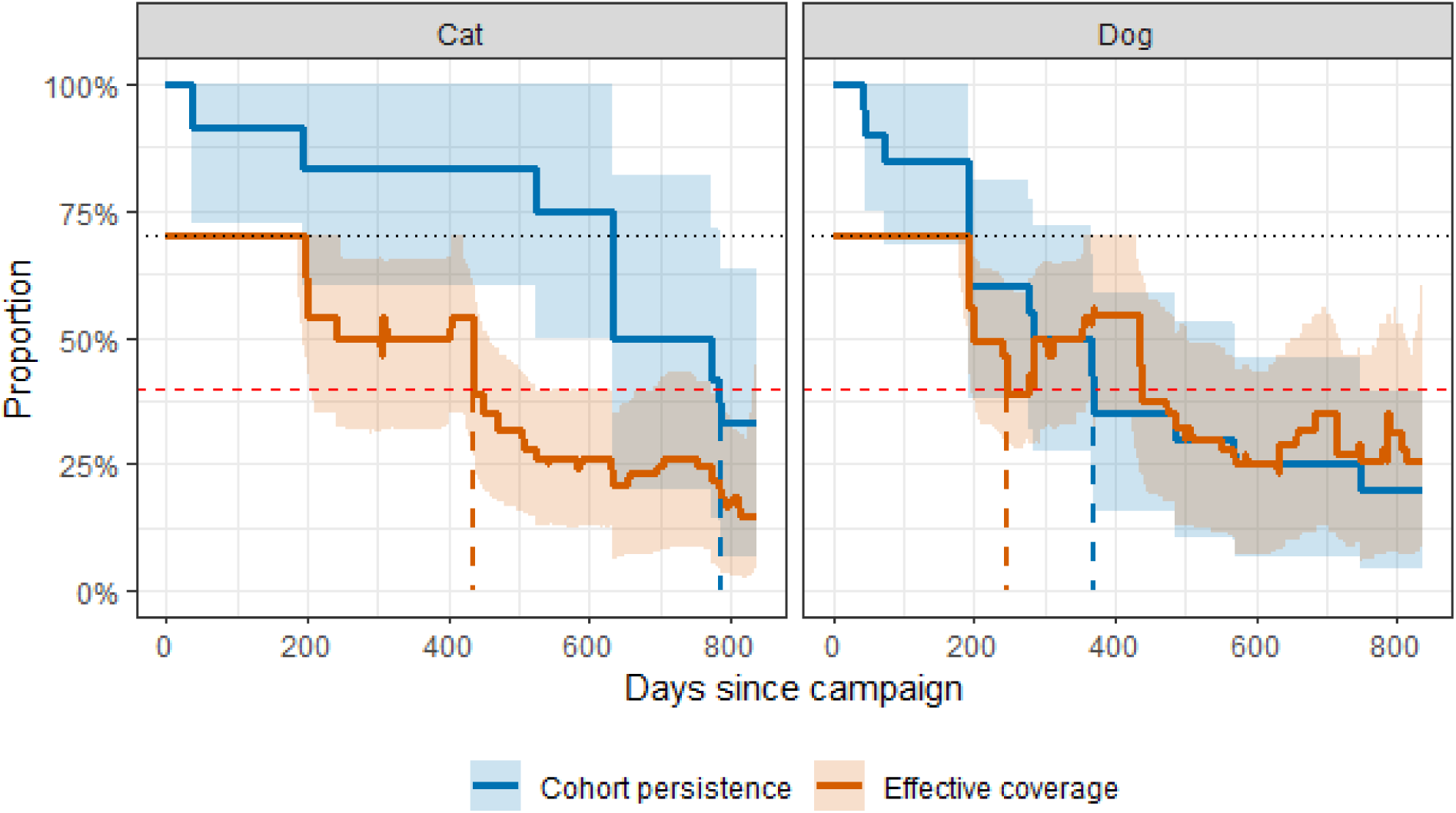
Observed erosion of herd immunity. Empirical comparison of cohort persistence and effective vaccination coverage over time. The blue line (Cohort persistence) represents the proportion of individuals present at the start of the study (t_0_) that remained on campus at each time point. The orange line (Effective coverage) represents the actual proportion of the total population that remains protected, calculated as the proportion of the total daily population (original cohort plus new recruits) that remained protected, assuming 70% initial vaccination coverage. The horizontal black dotted line is the initial coverage of 70% and the horizontal red dashed line indicates the critical herd immunity threshold (40%). Vertical dashed lines show when the critical threshold was passed. Divergence between the curves reflects immunity dilution due to recruitment of unvaccinated individuals.

While point estimates indicated that cohort persistence declined more slowly than effective vaccination coverage in both species, statistical confidence in this divergence varied. In cats, the 95% confidence intervals show a distinct separation between cohort persistence and effective coverage between approximately days 430 and 620, mathematically supporting the presence of immunity dilution during this specific window. In dogs, however, the confidence intervals overlapped almost entirely throughout the study period. This overlap indicates that while estimates suggest a buffering effect from declining abundance, the divergence in dogs is not statistically robust amidst the high baseline volatility of the population. The magnitude of dilution differed between species and was most pronounced in cats. By day 365, cohort persistence in cats remained high (83%), while effective vaccination coverage had declined to 50%, corresponding to a 33% divergence attributable to immunity dilution. This divergence isolates the loss of herd immunity due to recruitment of susceptible individuals, independent of mortality or emigration of vaccinated animals.

The mechanism of coverage loss differed fundamentally between species. Of the new cat recruits (n=58), 80.6% were juveniles (<6 months). Most of these juveniles were not born on campus (42% born, 58% abandoned). This suggests that on-campus reproduction was not necessarily the primary driver of dilution in the feline population. Conversely, new dog recruits (n=44) were predominantly adults (56.8%), consistent with transient or abandonment rather than on-site reproduction. Most of the new juvenile dogs were born on campus (63% born, 37% abandoned).

We quantified the attributable fraction of coverage loss for each species at Day 365. For cats, where cohort retention remained high (83%), the total coverage drop of 20 percentage points was driven by the recruitment of naïve individuals. Specifically, 40.5% of the reduction in herd immunity for cats was attributable to immunity dilution, with the remaining 59.5% caused by the loss of vaccinated residents. In contrast, estimates for dogs suggest that coverage loss was driven primarily by attrition. The dog population cohort retention rate reached 0.5 at Day 365. This metric remains mathematically distinct from the overall median persistence of 193 days derived from the Cox Proportional Hazards model, as cohort retention strictly tracks the proportion of the original vaccinated cohort remaining exactly one-year post-campaign. At Day 365, the observed effective coverage point estimate (53.8%) was higher than the theoretical coverage expected from retention alone, theoretically buffering the decline. However, the wide, overlapping confidence intervals for the dog cohort emphasize that these specific day-365 projections are subject to substantial variation. Rather than an exclusive failure mechanism, this wide variance highlights that dog herd immunity on campus is fundamentally unstable and highly sensitive to individual disappearance events.

## 4. Discussion

### 4.1. Summary of Main Findings

To our knowledge, this study provides the first direct comparison of demographic persistence and transient fractions in sympatric free-roaming dogs and cats. We identified an epidemiological divergence between the two species. While dog protection was lost primarily through the rapid exit of vaccinated individuals from the population (immunity attrition), cat protection was undermined chiefly by the continuous recruitment of naïve individuals (immunity dilution). This process driven by naive recruits created a 33% gap between individual cohort survival and actual population-level protection within one year. These findings suggest that standard annual vaccination pulses are mathematically insufficient for high-turnover urban animal populations. We discuss how effective One Health zoonotic control strategies must transition from static abundance-based targets to dynamic, species-specific intervention schedules.

### 4.2. Divergent Demographic Turnover and Epidemiological Failure Modes

#### 4.2.1 Population Fluidity and the Deception of Static Abundance

The free-roaming populations in this study demonstrated high fluidity, where the resident animals comprised a minority of the total population. Using species-specific resighting thresholds, 72% of dogs and 48% of cats were classified as transient. Population turnover, not size likely predicts the vulnerability of herd immunity on this and perhaps other university campuses. Our results indicate that a stable population provides better herd immunity than a larger, fluctuating population. Variation in observed abundances justifies the need for more effective environmental management rather than intermittent veterinary intervention alone. The free-roaming populations in this study demonstrated high fluidity. The functional resident population comprised a minority of the total identified cohort. Consequently, the study site appears to act as a transit node rather than a stable residential village. This turnover creates a high-risk sentinel site that constantly cycles naïve animals through the urban matrix.

The substantial individual variability observed in this study—visualized by the wide, heavily overlapping confidence intervals in vaccination coverage projections—demonstrates the fundamental danger of relying on static abundance estimates. In unmanaged urban areas, population turnover introduces extreme stochasticity; deaths, abandonments, or emigrations can drastically alter the actual immune fraction on any given day. This statistical volatility proves that the transition toward dynamic modeling remains a critical step for modern zoonotic disease control. Static annual targets cannot absorb this level of demographic instability, meaning intervention models must continuously update based on real-time residency and recruitment parameters to avoid silent failures in herd immunity.

#### 4.2.2 Species-Specific Failure Modes

The observed patterns and scenario results suggest that species-specific turnover rates drive divergent epidemiological failure modes. Our observed median persistence for dogs was approximately 6.4 months. This residency time is lower than that of owned roaming dogs in central Brazil (Belo et al., 2017), and falls within the high-turnover end of the demographic gradient described across South African and Indonesian study sites (Morters et al., 2014a, Morters et al., 2014b). In free-roaming dog populations in South Africa (Zenzele) and Indonesia (Bali) the majority of dogs had a life expectancy of at least 3 years, and 20% to 51.7% of starting cohorts remained present after 3 years (Morters et al., 2014b). While Morters et al. (2014b) established a paradigm of demographically stable free-roaming dog populations where annual vaccination campaigns likely remain effective (turnover was offset by relatively low emigration and immigration), our findings suggest that the Marco Zero campus instead functions as a high-turnover transient node.

For dogs at Marco Zero, the primary driver of vaccination failure was Immunity Attrition— defined as the loss of herd immunity due to the death or emigration of vaccinated individuals. In our simulated model, dog vaccination coverage collapsed below the 40% herd immunity threshold in only 160 days. The declining dog abundance created a concentration effect that temporarily buffered the decline in effective coverage due to Immunity Attrition. This “concentration effect” reflects a declining denominator, not genuine added protection. Observed dog sighting trends and low residency durations suggest the campus may act as a potential transient node in the broader urban matrix, facilitating a continuous circulation of naïve individuals, likely creating a high-risk cycle for zoonotic transmission.

Cat persistence was markedly longer than dog persistence. Even with relatively high individual persistence, immunity dilution was responsible for roughly 40% of the total erosion of herd immunity. We found that immunity dilution alone can create a substantial gap between cohort survival and cat population-level immunity. Cat sighting rates increased by 122% over the study period, consistent with continuous recruitment of naive individuals through reproduction and/or abandonment. For cats, juveniles comprised over 80% of new arrivals. Our decay model assumes demographic equilibrium, in which departing individuals are replaced at an equivalent rate. Comparison with observed abundance trends shows this assumption performed asymmetrically across species. For dogs, the declining population meant fewer naive animals entered the site than the model assumed. The modelled failure time (day 160) therefore represents a conservative safety margin. Observed effective coverage did not fall below the 40% threshold until day 248. For cats, the picture is more complex. The decay model predicted an apparent failure time of day 350. Empirical effective coverage did not however cross the 40% threshold until day 436 — later, not sooner, than modelled. This indicates the model’s built-in dilution assumption did not fully capture the actual balance of recruitment and persistence in cats. Nonetheless, the field data confirm that recruitment of naive juveniles remains a major driver of coverage loss in this species. While the decay model suggests an annual booster is sufficient for cats, the increasing relative abundance indicates a net influx of naive individuals. Therefore, the modeled failure time should be viewed as a best-case scenario. The actual herd immunity threshold may be breached much sooner than 350 days. This accelerated failure is driven by the continuous influx of susceptible kittens through abandonment and births.

### 4.3. Socio-Ecological Drivers and Targeted Intervention Strategies

#### 4.3.1 Socio-Ecological Drivers of Turnover

Addressing the “Dumping Ground” phenomenon is essential for stabilizing urban zoonotic disease control. High transient rates likely reflect frequent abandonment of pets on university grounds (Bicalho et al., 2024). This rapid turnover undermines the investment made in mass vaccination campaigns. If a relatively controlled environment cannot maintain herd immunity for six months, surrounding neighborhoods likely face even faster erosion. Data from these sentinel nodes should trigger broader municipal responses to prevent regional disease outbreaks. This turnover is not simply a feature of the surrounding urban landscape: it is compounded, and likely sustained, by the absence of any campus-level body responsible for admission control, sterilization referral, or adoption pathways at the study site — a governance gap documented at other Brazilian universities but rarely linked directly to its epidemiological consequences (Bicalho et al., 2024). Where no institutional mechanism exists to receive and act on that signal, as at present on the Marco Zero campus, the sentinel value of the site is being generated but not used.

#### 4.3.2 Targeted Management Logic

Educational campaigns are essential to shift the perception of the campus from a public shelter to a managed environment. The divergent dynamics identified in this study necessitate distinct management strategies for dogs and cats. The short-term dog strategy must focus on minimizing immunity attrition by reducing entry of transients and stabilizing the resident population. This requires enforcing control of illegal abandonment and implementing responsible guardianship campaigns in the nearby neighborhoods. These actions reduce the abandonment of unwanted animals on campus. As an annual vaccination cycle is virologically insufficient, a semi-annual pulse or continuous strategy is required.

Effective management within complex urban environments must transition from static abundance-based targets to dynamic, turnover-adjusted schedules. The cat strategy must prioritize the reduction of immunity dilution through sterilization. Vaccination alone is wasted if the population is continuously diluted by the birth and abandonment of susceptible kittens/juveniles. Sterilization has been proven to reduce population turnover and protect the long-term vaccination investment, but at the same time can drive compensatory immigration and increased survival in cats (Gerhold and Jessup, 2013, Gunther et al., 2022). Similar compensatory recruitment appears in other urban cat studies (Cove et al., 2023, Gunther et al., 2022). In some cases, Trap-Neuter-Vaccination-Return (TNVR) may therefore help maintain the herd fraction of immune individuals, but such actions are often impractical and likely to fail when conducted in isolation (Roebling et al., 2014, Read et al., 2020, Swarbrick and Rand, 2018). Sterilization and vaccination should therefore be treated as complementary tools, particularly given the growing recognition of cats as an under-addressed rabies host requiring dedicated management attention (Fehlner-Gardiner et al., 2024). In the case of the Marco Zero campus, life-extending veterinary care may effectively stabilize dog coverage but will fail to protect cats without concurrent controls on reproduction. These species-specific failure modes require different intervention logics. Disappearance of dogs represents an absolute loss from the study area. Increasing cat abundances suggest compensatory recruitment into a resource-rich niche. Individual-based photographic tracking and Cox models are necessary to parameterize these distinct processes, particularly in complex urban settings where populations fluctuate dynamically. This monitoring should form part of integrated approaches that include formal governance and inclusion of local communities to facilitate adoption and reduce abandonment (Swarbrick and Rand, 2018, Nayeri et al., 2025).

### 4.4. Institutional Frameworks and the Sentinel Site Model for One Health

University campuses are semi-permeable sites well suited to sentinel surveillance (Guillot et al., 2022). Beyond its immediate demographic findings, this study illustrates how a spatially bounded, non-invasively monitored site can serve as a sentinel for population processes that are difficult to observe at city-wide scale. Our low-cost approach could work at other institutions without heavy infrastructure. This value, however, is contingent on institutional uptake: a sentinel site without a receiving governance structure generates surveillance data that cannot be translated into sustained management action, however sophisticated the underlying method. Globally universities have institutionalized comparable free-roaming populations through designated management committees and trap-neuter-vaccinate-return programmes (Phiriyaphokhai et al., 2025, Swarbrick and Rand, 2018); at present, UNIFAP has neither. Our research demonstrates the potential of the Marco Zero campus as a sentinel site for evaluating population dynamics. We recommend formalizing the presence of resident community animals within institutional sustainability plans. Adopting a One Health approach and acknowledging these animals as biological indicators of environmental health legitimizes management efforts (Schneider et al., 2023). Our monitoring framework provides the longitudinal data necessary to detect changes in turnover without requiring expensive equipment. Strategic partnerships between universities and municipal zoonosis control centers are vital. The university can provide high-resolution surveillance data to identify stable residents versus transient populations. In exchange, municipal authorities could provide vaccines and castration slots. Beyond the health benefits of vaccination and castrations, this integration is likely to also increase chances of adoption, which remains one of the most challenging aspects of control programs. At a minimum, delivering on this requires naming a responsible unit (for example, a One Health or animal-welfare working group with student, veterinary, and administrative representation), assigning a budget line, and adopting transparent governance policies.

#### 4.4.1 Scaling the Sentinel Site Model

Institutional partnerships create a cost-effective, evidence-based feedback loop for urban disease management. Such collaborations ensure that zoonotic disease control strategies are adjusted in real-time based on measured rates of dilution. Traditional census counts could be strengthened by demographic surveillance systems, as used elsewhere for dogs (Conan et al., 2015). This tracks tenure and turnover, not just abundances. Vaccination targets should be dynamic and adjusted in real-time. If turnover increases due to post-holiday abandonment spikes, vaccination efforts must increase proportionally. Data from these sentinel sites should feed into broader urban health planning. A change in turnover on campus can be used to trigger responses in waste management and public health messaging in neighboring areas (Bastos et al., 2021, Silva et al., 2024). This mirrors Brazil’s own history of One Health rabies control (Schneider et al., 2023).

Furthermore, this study demonstrates that robust epidemiological science can be generated through methodological ingenuity rather than infrastructural abundance. By pairing low-cost, non-invasive photographic mark-resight surveys (using mobile phones) with robust statistical techniques like non-parametric bootstrapped survival analyses, we successfully extracted high-resolution turnover parameters. This approach bypasses the need for expensive GPS collaring or invasive trapping, proving that resource-limited municipalities and universities can still generate the demographic data required to parameterize dynamic One Health models.

The extreme epidemiological volatility observed at the Marco Zero campus—characterized by high turnover and wide confidence intervals is a direct symptom of an unmanaged ‘dumping ground’ dynamic. Therefore, the absence of institutional oversight at this site is not merely a study caveat, but an actionable finding. A sentinel site without a receiving governance structure generates vital early-warning data that ultimately goes to waste. To fix this, universities must establish a minimum viable governance structure: a formalized One Health or Animal Welfare Committee comprising university administration, veterinary/biological sciences faculty, and student representatives, supported by a dedicated budget line. This body must help inform campus admission controls (anti-abandonment policies), manage formalized adoption pathways to stabilize the resident population, and coordinate directly with municipal zoonosis centers to exchange surveillance data for vaccination and sterilization resources. Until such governance is implemented, campus populations will remain a self-renewing source of zoonotic risk rather than a controlled sentinel node.

### 4.5 Caveats

Our focus is to inform preemptive prevention, not outbreak control, which needs different data including population numbers and movement patterns (Cunha Silva et al., 2025). Our single-site study thresholds are benchmarks, not universal targets. Extending this monitoring to more sentinel sites would show whether these patterns hold more broadly. While abundance and turnover dynamics may vary, our monitoring framework is likely to be widely applicable across university campus sentinel sites, especially those with biological and/or environmental science courses. Sampling covered only the dry season and term time, so seasonal abandonment spikes could not be assessed. Our “loss” event conflates confirmed death with unconfirmed emigration, which have different epidemiological meanings. As no animals were vaccinated all coverage figures exclude vaccine-titre waning, which would probably add to, not offset, the demographic losses modelled here. Our estimated failure times should be treated as upper bounds on true protection durability, not as forecasts of an implemented campaign. Future work should prioritize multi-site, multi-season sentinel networks, plus real vaccination and serological monitoring.

### 4.6 Conclusion

Demographic turnover was the primary barrier to maintaining herd immunity in an urban sentinel site. We have shown that the mechanisms of immunity loss are species-specific and demand divergent management logic. Effective One Health strategies must transition from static abundance targets to dynamic, turnover-adjusted vaccination schedules. By addressing Immunity Attrition in dogs and immunity dilution in cats, cities can better manage zoonotic risks. Integrating field data into broader urban health planning will allow for more resilient disease control interventions. Extending this monitoring to more sentinel sites would show whether these patterns hold more broadly. More broadly, the epidemiological value of a sentinel site is limited by its governance. Without a designated institutional structure to convert turnover data into vaccination, sterilization, or adoption action, even a well-monitored campus population will continue to function as a self-renewing source of zoonotic risk rather than a controlled one.

## Acknowledgements

We are grateful to the numerous undergraduate students who participated in the monitoring activities over the years. In particular we thank undergraduate students Bruno, Carina, Halbertt, Sâmela and Vinicius for their dedication.

