## Supplemental material for "Demographic turnover undermines herd immunity in sympatric free-roaming dogs and cats: implications for zoonotic disease control in urban sentinel sites"

### S1 GAM model predictions

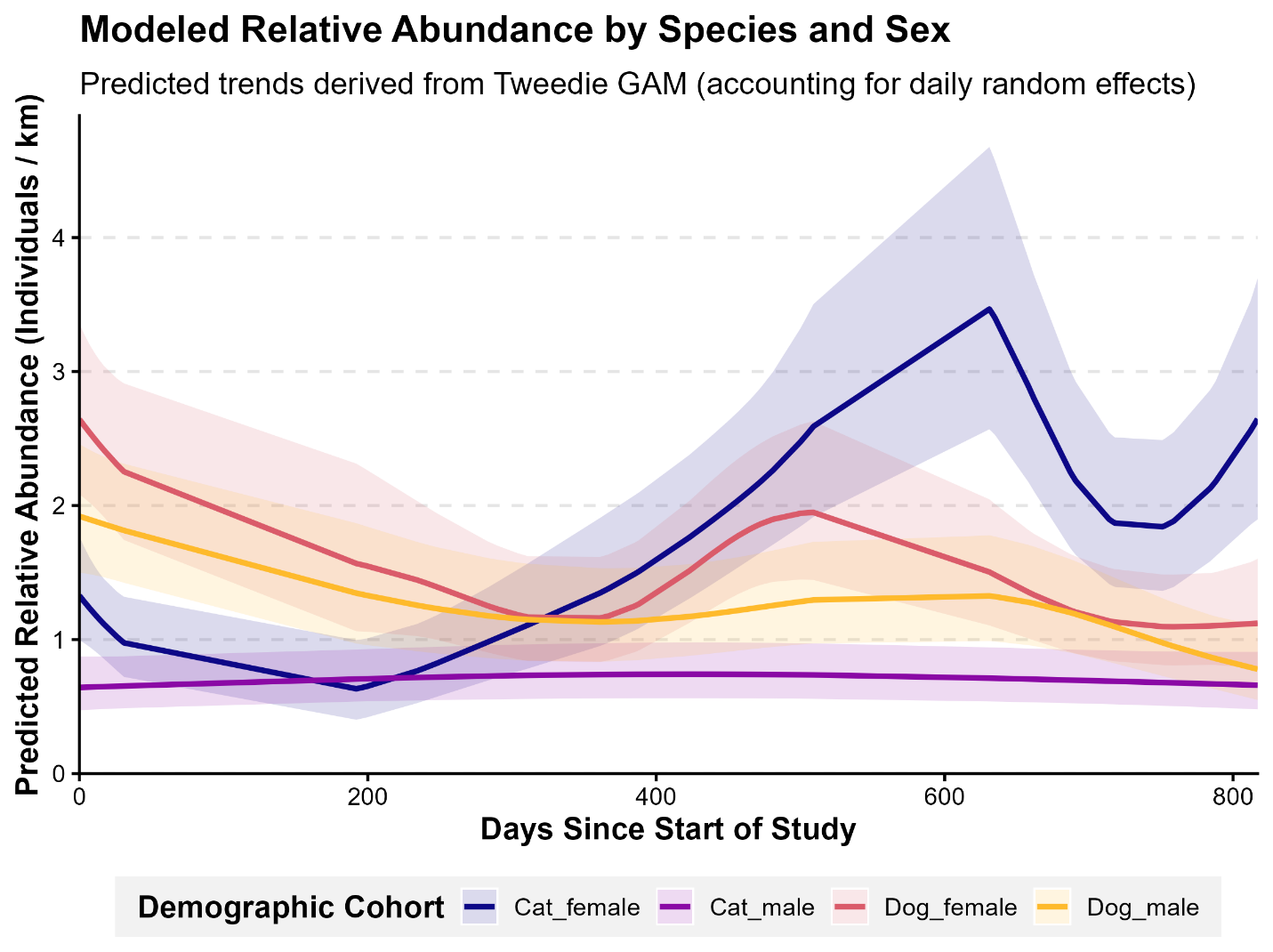

### S2 Accumulation of individuals

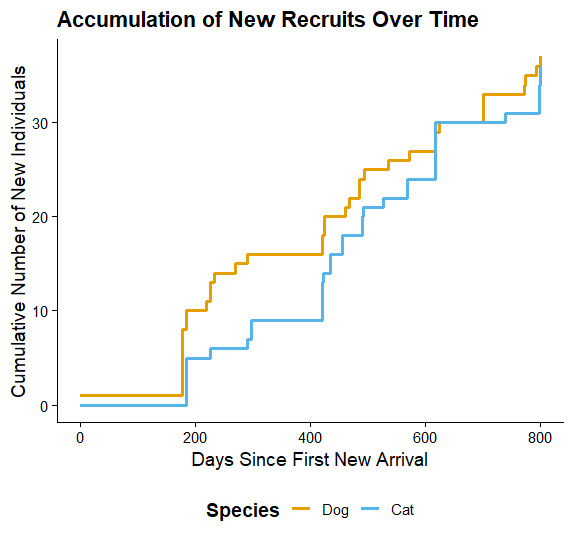

### S3 Full Cox Proportional Hazard

Results for model including adopted individuals.

Analysis revealed no significant effect of sex/species on persistence. Cats had a 30% lower hazard (risk of leaving/dying) compared to dogs, holding sex constant. This suggests cats tend to persist on campus longer than dogs. Males have approx. 46% higher hazard (risk of leaving/dying) compared to females, holding species constant. This suggests males tend to disappear or die faster than females. Even though the results were not statistically significant, the effect sizes are ecologically notable (a 30% reduction and 46% increase in risk are large magnitudes).

The estimated median residency time for cats was nearly double that of dogs (600 and 342 days for cats and dogs respectively). This means that, adjusting for covariates, an individual cat has a 50% probability of surviving past day 600. There was variation between sexes of both cats and dogs, with males remaining for less time compared with females (Figure 4). Under the mortality conditions observed in this study, we expect 50% of the initial cohort to have left the campus by time $X$.

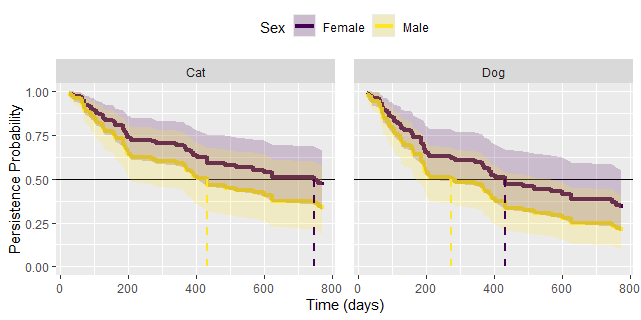

Figure S3. Persistence Probability. Results of Cox Proportional Hazard model showing the probability of cats and dogs remaining on campus over time. Solid lines represent the estimated probability over time for males and females. Shaded regions indicate 95% confidence intervals. Dashed vertical lines mark the median time for each group.

### S4 Turnover rate in vaccination coverage decay

To support the comparison made in the Discussion — that turnover rate, not population size, determines the durability of herd immunity — we used the vaccination-coverage decay equation from Methods 2.3.3 $V_{t}= V_{0}. e^{-\lambda t}$ to project two hypothetical population trajectories forward 365 days. Because this equation contains no population-size term, coverage decay is mathematically independent of N; only the turnover-driven decay constant (λ) determines the trajectory. A "closed" trajectory (λ = 0) represents a population with no exits and no new recruits; a "fluctuating" trajectory uses the empirical dog population half-life (T½ = 193 days, Results 3.3) as a realistic turnover rate. We ran two paired comparisons: as stated in the Discussion (50 dogs, closed, versus 30 dogs, fluctuating), and with sizes reversed (30 dogs, closed, versus 50 dogs, fluctuating), to test whether the pattern held even when population size favored the fluctuating group. Under the fluctuating trajectory, coverage fell below the 40% herd-immunity threshold by day 156, irrespective of whether N was 30 or 50. In the reversed comparison, the smaller closed population (30 dogs) overtook the larger fluctuating population (50 dogs) in absolute numbers of protected animals by day 142, and remained ahead for the remainder of the study period.

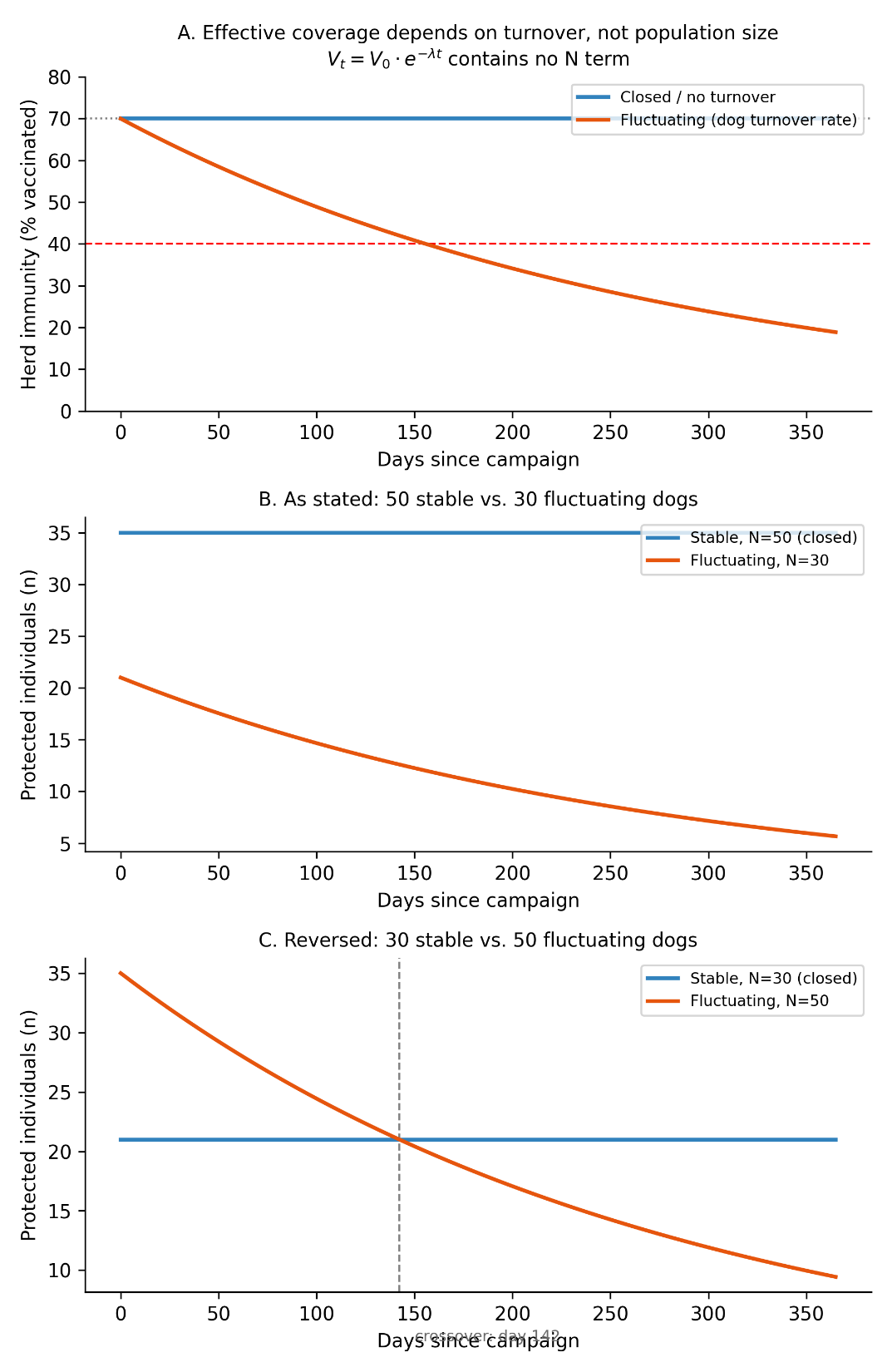

**Figure S4. Turnover rate, not population size, determines the durability of effective vaccination coverage.** Hypothetical illustration using the coverage-decay equation from Methods 2.3.3, parameterised with the empirical dog turnover rate (T½ = 193 days) and a closed-cohort boundary case (no turnover). (A) Percentage coverage over time for a closed population (flat, never declines) versus a fluctuating population (declines below the 40% threshold, red dashed line, at day 156); this curve is identical regardless of population size. (B) Absolute number of protected individuals over time for the comparison as stated in the Discussion: 50 dogs, closed, versus 30 dogs, fluctuating. (C) The same comparison with sizes reversed: 30 dogs, closed, versus 50 dogs, fluctuating. The grey dashed line marks day 142, when the smaller closed population overtakes the larger fluctuating population in absolute protected numbers.

### S5 Key Data & Definitions

**Table S5.1: Comparative Demographic Metrics (UNIFAP)**

| **Metric** | **Dogs (C. l. familiaris)** | **Cats (F. catus)** |
| --- | --- | --- |
| **Sample Size** | N=72 | N=75 |
| **Transient Rate** | 72% | 48% |
| **Median Persistence** | 193 Days | 432 Days |
| **Hazard Ratio (HR)** | 1.0 (Ref) | 0.56 ($p=0.029$) |
| **Herd Immunity Failure (<40%)** | Day 160 | Day 350 |
| **Demographic Mechanism** | High Attrition (Exit) | High Dilution (Entry) |

**Table S5.2: Glossary of Used Terms**

| **Term** | **Definition** | **Context Usage** |
| --- | --- | --- |
| **Immunity Attrition** | Loss of herd immunity due to the death or emigration of vaccinated individuals. | "Rapid immunity attrition in dogs renders annual vaccines ineffective." |
| **Immunity Dilution** | Reduction in the proportion of immune individuals due to the recruitment (birth/immigration) of naïve animals. The epidemiological term for when the proportion of immune individuals drops because new, susceptible individuals (unvaccinated immigrants or newborns) enter the population.  • Why it fits: It perfectly describes your "sink" scenario where the population size might stay stable, but the "concentration" of immunity is diluted by the rapid replacement of animals. | "Cat vaccination coverage was eroded by immunity dilution despite high survival."  "The high rate of population turnover results in rapid immunity dilution of herd immunity, rendering annual vaccination pulses ineffective." |
| **Broader Term: "Vaccination Coverage Decay"** | This is the standard term used in quantitative modeling (e.g., Hampson et al., Cleaveland et al.) to describe the decline in the percentage of vaccinated animals over time.   - **Why it fits:** It is neutral and mathematically rigorous. It encompasses both the death of vaccinated animals and the birth/immigration of susceptible ones. | *"We modeled the* ***vaccination coverage decay*** *to determine the critical time points at which herd immunity fell below the 40% threshold."* |
| **The Mechanism-Specific Term: "Immunity Attrition"** | Used to highlight the *loss* of the vaccinated animals themselves (death or emigration).   - **Why it fits:** It emphasizes the exit of protected animals rather than the entry of new ones. | *"High mortality rates among free-roaming dogs drive rapid* ***immunity attrition****, necessitating continuous rather than pulsed intervention strategies."* |
| **Sympatric Divergence** | The phenomenon where co-occurring species exhibit differing demographic or behavioral responses to the same environment. | "Sympatric divergence in turnover necessitates differential management." |
| **Sentinel Site** | A specific location monitored to detect changes in the broader epidemiological landscape. | "The campus acts as a sentinel site for urban rabies risk." |
